# Comprehensive nucleosome mapping of the human genome in cancer progression

**DOI:** 10.1101/021618

**Authors:** Brooke R. Druliner, Daniel Vera, Ruth Johnson, Xiaoyang Ruan, Lynne M. Apone, Eileen T. Dimalanta, Fiona J. Stewart, Lisa Boardman, Jonathan H. Dennis

**Affiliations:** Department of Biological Science, Florida State University, Tallahassee, Florida, United States of America; Department of Laboratory Medicine and Experimental Pathology, Mayo Clinic, Rochester, Minnesota, United States of America; Biomedical Statistics and Informatics, Department of Health Sciences Research, Mayo Clinic College of Medicine, Rochester, Minnesota, United States of America; New England Biolabs Inc., Ipswich, Massachusetts, United States of America; Division of Gastroenterology and Hepatology, Department of Internal Medicine, Mayo Clinic, Rochester, Minnesota, United States of America

**Keywords:** cancer, chromatin, nucleosome, MNase, whole genome, Next Generation Sequencing, lung adenocarcinoma, colorectal carcinoma, mTSS-seq

## Abstract

Altered chromatin structure is a hallmark of cancer, and inappropriate regulation of chromatin structure may represent the origin of transformation. Important studies have mapped human nucleosome distributions genome wide, but the role of chromatin structure in cancer progression has not been addressed. We developed a MNase-Transcription Start Site Sequence Capture method (mTSS-seq) to map the nucleosome distribution at human transcription start sites genome-wide in primary human lung and colon adenocarcinoma tissue. Here, we confirm that nucleosome redistribution is an early, widespread event in lung (LAC) and colon (CRC) adenocarcinoma. These altered nucleosome architectures are consistent between LAC and CRC patient samples indicating that they may serve as important early adenocarcinoma markers. We demonstrate that the nucleosome alterations are driven by the underlying DNA sequence and potentiate transcription factor binding. We conclude that DNA-directed nucleosome redistributions are widespread early in cancer progression. We have proposed an entirely new hierarchical model for chromatin-mediated genome regulation.

**Significance:** This is the first report of human nucleosome distribution in cancer progression using sequence capture and HiSeq. We show in lung and colorectal adenocarcinoma patients that nucleosome distribution is a widespread, early response driven by genetically-encoded signals and potentiate regulatory factor binding. We present a model for chromatin-based hierarchical regulation.

## INTRODUCTION

Despite the central role of chromatin as the ultimate substrate for all nuclear events, the structure of chromatin remains poorly characterized. The human genome is packaged into chromatin, whose fundamental subunit is ∼147bp of DNA wrapped around a histone octamer to form the nucleosome ^1^. The location and density of nucleosomes with respect to the underlying DNA sequence is an important factor in determining access to the genome for DNA-templated processes^2^^-^^4^. Little is known regarding the precise role of nucleosome distribution in these processes, because there have been relatively few studies measuring the distribution of nucleosomes across the genome in multiple cell types and physiological contexts.

Genome-wide nucleosome distribution information is critically important for understanding genomic processes, yet this information is lacking for a variety of human cell states. Genome-wide measurements of the locations of genome binding factors by Chromatin immunoprecipitation (ChIP), polymorphisms by exome sequencing, or DNA methylation by bisulfite conversion, have become routine and robust assays of genomic structure and organization. A literature search on any of these assays returns thousands of results, while searches on “nucleosome distribution” returns an order of magnitude fewer results. Only a handful of seminal papers have measured genome wide human nucleosome positions in a limited number (1-2) of cell states ^5^^-^^8^. To address this shortcoming, we have developed a robust, cost-effective sequencing-based nucleosome distribution mapping platform to analyze chromatin structure at the transcription start site (TSS) of ∼22,000 open reading frames in the human genome. We have applied this approach to study the nucleosome distribution in primary patient tumor samples representing multiple stages and grades of both lung adenocarcinoma (LAC) and colorectal cancer (CRC).

A complete understanding of the distribution of nucleosomes across the genome in cancer is currently lacking, yet it is critically important for understanding cancer etiology in basic biological and clinical contexts. We have previously shown extensive nucleosome distribution changes at a subset of genes in patients with low-grade LAC ^9^.

Mononucleosomally protected DNA was isolated from patient derived primary LAC tissue and used to query high-resolution tiling microarrays. These microarrays were custom-designed to measure nucleosome distribution changes at the 2000 bp surrounding the TSS of ∼900 cancer- and immunity-related genes. Those studies were limited in the breadth of loci studied by the number and density of probes that it was possible to print on the microarray. In the present report, we have redesigned the original experimental approach for the targeted sequencing of TSSs by paired-end sequencing.

We have developed a solution-based sequence capture method enabling the enrichment of the 2000 bp surrounding the TSS of ∼22,000 open reading frames in the human genome. Due to the importance of promoter composition in gene regulation, we designed the method to map nucleosomes at the regions surrounding the TSS. This capture method reduces the sequence space of the human genome from 3.4 Gb in total to ∼50Mb of TSSs, a 98.5% reduction. This enrichment is analogous to that achieved for well-documented exome sequencing experiments ^10^. Using this targeted enrichment of mononucleosomally-protected DNA, which we call mTSS-seq (MNase-protected DNA, transcription start site capture-sequencing), we were able to achieve high enough sequencing coverage to determine individual nucleosome positions, at an average of ∼100 reads per nucleosome, exceeding the necessary coverage for high-resolution nucleosome position mapping ^5^^,^^7^^,^^11^^,^^12^.

This technique represents a unique source of nucleosome distribution information at the TSS, and has not been previously executed on a genome-wide scale. The relative enrichment or reduction of sequences from this assay allows us to determine changes in nucleosome distribution among a variety of sample types. This approach offers several clear advantages. Our approach measures nucleosome distribution at all TSS in the human genome. The targeted enrichment is a cost-effective approach to whole genome studies and allows for comprehensive nucleosome distribution mapping to be completed on several samples. This nuclease protection assay is highly relevant to diffusible molecules such as transcription factors and the paired end sequencing approach provides information on protected fragment size. We can therefore use this assay to analyze subnucleosomal-sized fragments for an additional layer of genomic regulatory information ^11^^,^^13^. Using our newly developed mTSS-seq approach, we have mapped nucleosome distribution with unprecedented breadth and depth in human cancer patient samples.

## RESULTS

### Development of a solution-based TSS-enrichment sequence capture method for mononucleosome DNA from primary patient tissue

In this study, we measured genome-wide chromatin structure in primary patient tumors using a newly developed approach that we present here for the first time. At the outset, we used matched tumor and normal tissue from grade one and three LAC patients on which we have previously reported ^9^. The workflow is shown in Fig. 1A where, following digestion with MNase, mononucleosomal DNA was isolated (all material below ∼150bp was excised from a 2% agarose gel). In many cases we used the exact mononucleosomally protected DNA sample prepared for the original report on these patients (Supplementary Table 1 – MNase preparation column). This MNase digested, mononucleosome DNA material was used to prepare multiplexed libraries. Notably, selection of material from the nucleosomal ladder that was 150bp and lower allowed for analyses of subnucleosomal fragments derived from other non-histone DNA-binding proteins such as transcription factors. We efficiently tracked addition of adaptor and barcode ligated material at every step of the library preparation and accurately quantified the exact number of molecules adjusted for size to be sequenced. This ensured that the majority of the reads we obtained would strictly give nucleosome and subnucleosome information.

**Figure 1.**
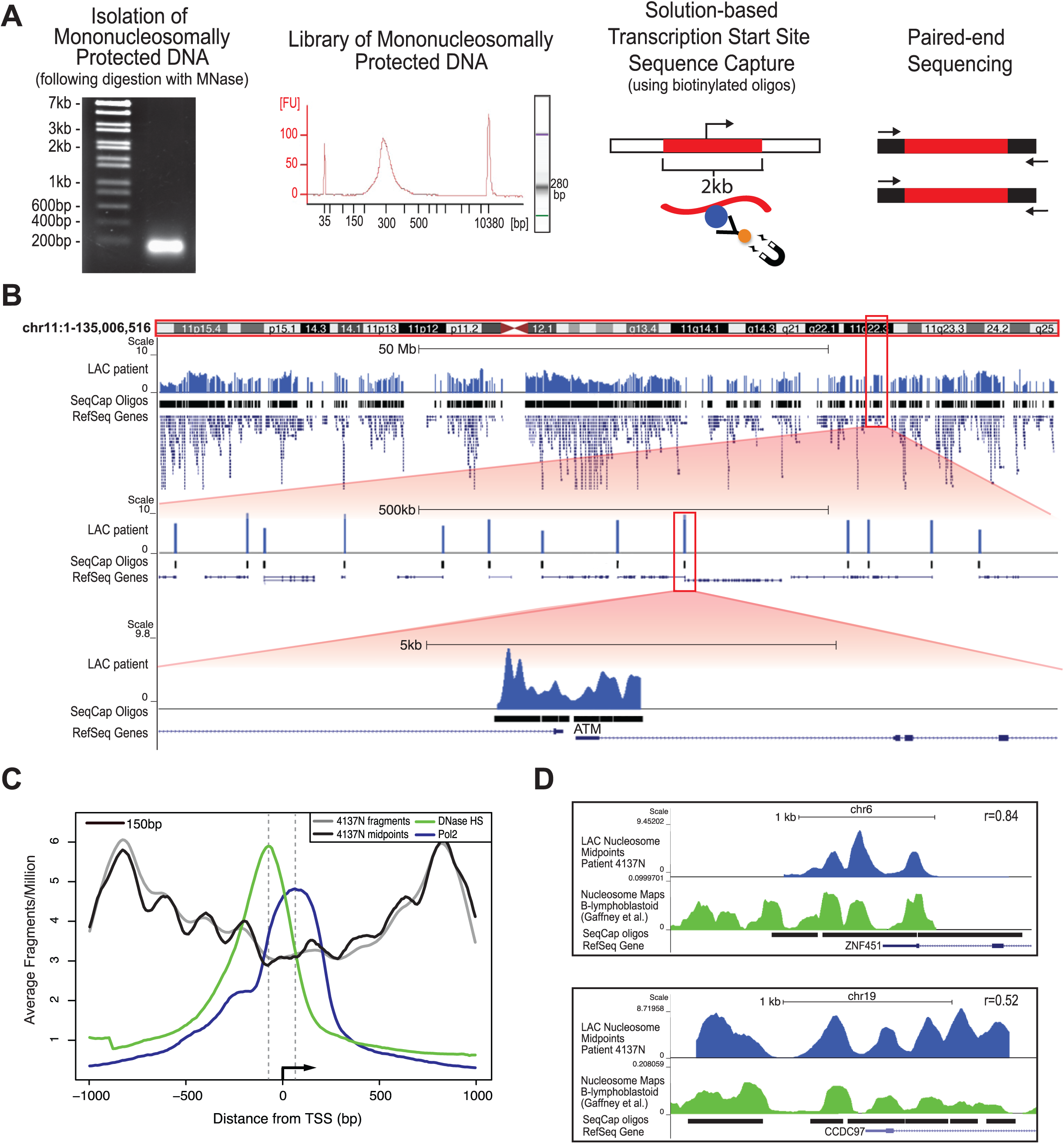
**The newly developed mTSS-Capture method combined with paired-end sequencing maps genome-wide nucleosome distribution in primary patient samples and identifies *bona fide* nucleosome characteristics, concordant with other human nucleosome mapping studies.** (A) Work-flow of the mTSS-seq method. Following MNase digestion using a titration of MNase, populations of mononucleosomally protected DNA and subnucleosomal fragments are isolated, and prepared as libraries for Illumina sequencing. Solution-based sequence capture is performed using biotinylated oligos, enabling the enrichment of fragments within 2kb of each transcription start site in the human genome. Paired-end 50bp sequencing was then performed on each index. (B) Alignment of the mTSS-seq midpoints to the human genome using the UCSC genome browser for LAC patient #4137 Normal tissue is shown for chr11, hg19 (<u>http://genome.ucsc.edu</u>). Zooming in twice at 100X allows for further visualization of the sequence capture oligos surrounding the TSS in a 500kb and a 5kb region showing the ATM locus. (C) Averaged, normalized reads per million (y-axis) from mTSS-seq plotted as fragments (gray) and midpoints (black), centered on and surrounding 2kb of the TSS for ∼22,000 open reading frames in hg19 (x-axis). DNase I-hypersensitivity (GSM736580; green) and RNA polymerase II from ChIP-seq (GSM935299; blue) data from A549 cells are shown. (D) LAC patient 4137 Normal nucleosomal midpoints (blue track) were plotted in the UCSC genome browser against the published human lymphocyte nucleosome distribution maps by Gaffney et. al. (green track) for the ZNF451 and CCDC97 loci. Sequence capture oligos and corresponding RefSeq gene models are shown for each locus. Correlations are shown for ZNF451 and CCDC87, respectively.

Following preparation of the libraries, we used our custom-designed solution-based sequence capture to select the 2000bp surrounding the TSS of ∼22,000 human open reading frames, allowing us to capture nucleosomes covering ∼48Mb of the human genome. Prior to performing paired-end sequencing on the captured material, we quantified the enrichment of our sequence capture by qPCR using specific primers to regions on-target and off-target from the capture (Supplementary Table 2). In the captured libraries, the difference between the on-target (enriched) and off-target (depleted) C_T_ values differed at an average of 9 cycles, a minimum of 100-fold enrichment (Supplementary Figure 1A). Following sequencing, we aligned the paired-end reads to the human genome and determined the size of each fragment from the separation of the paired ends (Supplementary Figure 1B). The majority of the sequenced fragments were within the 75-200bp size range, showing that the range of sizes across samples was relatively consistent. Figure 1B shows the frequency of inferred nucleosome midpoints in genome traces across chromosome 11, and zoomed in twice at 100× to eventually show a single locus (TSS of the ATM gene) and the resulting nucleosome distribution map (Fig. 1B). The sequence capture oligos used to capture the 2kb surrounding the TSS are shown in this view, along with the data corresponding to the targeted regions, which in every case made up over 90% of the total sequencing reads (Supplementary Table 3).

### Paired-end reads generated by mTSS-Seq yield typical nucleosome characteristics, and are concordant with previous reports in the literature

To validate the use of mTSS-seq to accurately map nucleosome distribution we identified typical nucleosome characteristics in our data, and compared our data to other published human nucleosome mapping studies. To determine whether our data contained typical nucleosome properties we plotted both the average nucleosome distribution for all TSSs in the genome and determined dinucleotide frequencies. Nucleosome organization averaged around the TSS of ∼22,000 human genes shows a canonical structure with phased nucleosomes centered on a nucleosome depleted region ^7^. We determined the average nucleosome organization at the TSS for our data by aligning all TSSs and plotting the corresponding sequence fragment midpoints for the 2kb surrounding the TSS (Fig. 1C). Our mTSS-seq data recapitulate the pattern of other studies that plotted average nucleosome occupancy at the TSS in humans, where there is a NDR (nucleosome-depleted region) surrounding and immediately downstream of the TSS, with well-positioned nucleosomes flanking the NDR ^7^. Additionally, we demonstrated that the NDR directly upstream of the TSS overlaps with a peak in genomic DNase I hypersensitivity, and a ChIP peak for RNA polymerase II is seen just downstream of the TSS within the gene body, as anticipated ^7,14-17^. We further validated our technology with a sequence analysis of the nucleosome-sized fragments generated by mTSS-seq.

A major determinant of the ability of DNA to conform to the histone octamer into a nucleosome is the specific patterns of dinucleotides^18^. Specifically periodic AA distributions occur in sequences higher than expected, and are thought to be responsible for genome organization into nucleosomes ^19^^-^^22^. The periodic occurrence of A/T containing dinucleotides at ∼10 bp intervals was calculated from first principles and verified in several subsequent studies 23-26. When we examined the dinucleotide frequency of 150 bp fragments, we found the acknowledged 10bp periodicity for A/T containing dinucleotides, comparable to the frequency patterns identified in other human studies (Supplementary Figure 1C) ^5^^,^^26^^,^^27^. These results affirm that the high resolution maps provided by mTSS-seq are consistent with the major qualitative features of nucleosome distribution at the TSS, and with the sequence composition of bona fide nucleosomal particles.

We next wanted to verify that our mTSS-seq data agreed with precedent human nucleosome mapping studies at specific loci. This comparison is particularly important, as averages and qualitative measures of general nucleosome distributions are not necessarily sufficient to make claims about nucleosome organizations at specific loci. We compared the nucleosome distribution data of normal lung epithelial patient tissue from our study (patient #4137N) to data from a human lymphoblastoid cell line, and found a positive global correlation of 0.37 ^5^. We have shown two representative examples at the loci ZNF451 (r=0.84) and CCDC97 (r=0.52), demonstrating the similarity between our mTSS-seq derived data and the lymphoblastoid cell line data (Fig. 1D). We next validated the mTSS-seq data by comparison to our previously published microarray-based study ^9^. We have been able to detect changes in nucleosome distribution between normal and tumor tissue in the early LAC patients, and these changes are consistent with those observed by microarray (Supplementary Figure 1D). Our comprehensive mTSS-seq approach therefore allows the generation of nucleosome distribution maps and measurement of changes between samples with similar accuracy to previously published studies, with an increased breadth, querying all TSSs of the human genome.

### mTSS-Seq identifies specific nucleosome architectures and genome-wide nucleosome distribution alterations in the progression of LAC

Our previous study demonstrated that nucleosome redistributions occurred at 50% of the ∼900 TSS studied. It was important to determine whether the widespread nature of these changes was limited to the loci studied in the previous investigation, or whether these changes were part of a larger genome wide nucleosomal reorganization. To investigate genome-wide changes in nucleosome distribution at the TSS, we first calculated the difference between the normal and tumor datasets for each patient. Sorted difference maps of these data show that the grade one nucleosome distribution differences are widespread and dispersed throughout the genome, while these differences are greatly diminished in both grade three patients (Fig. 2A). Additionally, the difference maps show that the widespread changes observed exclusively in the grade one patients are associated with a lower occupancy in the tumor as compared to normal. These results demonstrate that the nucleosome distribution changes exclusive to the grade one tumors are widespread and shows lower occupancy at the TSS across the entire human genome.

**Figure 2.**
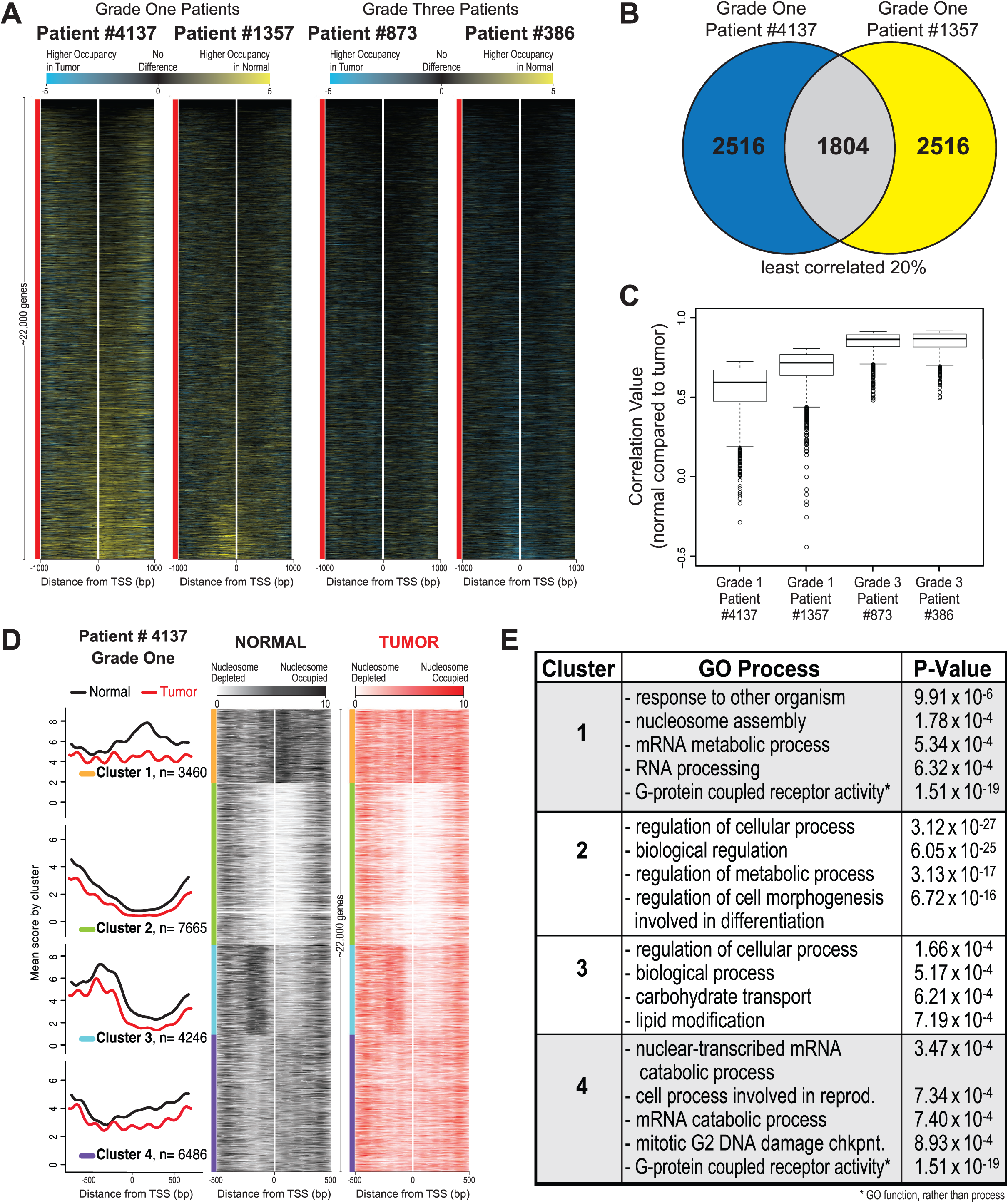
**Figure 2 Widespread nucleosome distribution changes are common between grade one patients and have specific nucleosome architectures that are enriched for specific GO processes.** (A) Heatmaps representing Normal minus Tumor differences are shown for each patient at each TSS in the human genome (∼22,000 genes). Loci are sorted on the mean difference value across 2000 bp surrounding the TSS (white line) for all genes, ordered on the basis of each patient’s corresponding normal data. The two grade one patients are on the left and the two grade three patients are on the right. Black represents areas with few differences between normal and tumor, yellow (positive values) indicates higher nucleosome occupancy in the normal and blue (negative values) indicates higher nucleosome occupancy in the tumor. (B) Correlation values were calculated between normal and tumor for each grade one patient, and the genes in the lowest 20% (indicating change between normal and tumor) were selected (∼4,300 genes each). The overlap of genes determined as changed for each grade one patients is shown, 1,804 genes. (C) Correlation values between normal vs. tumor for each of the common 1,804 genes are plotted as boxplots for each patient, showing a range of lower correlation values in both the grade one patients (left two boxplots) as compared to the range of higher correlation values in the grade three patients (right two boxplots). (D) The average nucleosome occupancy and corresponding heatmaps are shown for the Normal (black) and Tumor (red) data for grade one patient #4137, generated using k-means clustering with k=4 centered on the 1000bp surrounding the TSS for every human gene. The four clusters contain 3,460, 7,665, 4,246 and 6,486 genes, respectively. In the average plots, the y-axis is the mean score for both normal (black) and tumor (red) data, for each cluster. The x-axis is the genomic position for both the average plots and heatmaps. In the heatmaps, white represents nucleosome depletion and black (normal) or red (tumor) represents nucleosome occupancy. We determined that the majority of the 1,804 genes identified in part B of this figure belonged to clusters 1 and 4. (E) The enrichment of genes in each cluster for a ontologic process was calculated. The four processes most overrepresented by genes are shown for each cluster with corresponding *p*-value.

In order to determine if the genome wide nucleosome distribution changes in grade one tumors were similar between patients, we quantified the overlap in genes with nucleosome distribution alterations between the grade one patients. We first calculated the correlation between normal and tumor for each grade one patient for every gene, and then identified overlapping genes in the least correlated 20% (∼4,300 genes). We found that 1,804 genes with the greatest degree of change between normal and tumor overlapped between the grade one patients (Fig. 2B). Additionally, the grade one patients showed a broad range of low correlation values (ranging from -0.5-0.8), whereas the grade three patients had a smaller range of values (ranging from 0.5-0.9) (Fig. 2C). The significant degree of overlap in nucleosome distribution changes suggests a concerted set of nucleosome distribution changes for these loci in early adenocarcinoma.

To test for nucleosome distribution organizations in the early tumors that might indicate shared chromatin structural events in early LAC, we categorized nucleosome profiles surrounding each TSS in the genome. We used *k*-means to align and cluster all genes based on nucleosome occupancy for a patient tumor and matched normal tissue (Fig. 2D). Since it was previously determined that four significantly distinct clusters defined nucleosome architectures, we used *k*=4 for *k*-means clustering of our data, and grouped the profiles in a window of 1000bp surrounding the TSS of the entire genome ^12^. We found that decreasing the number of clusters combined clusters 1 and 4, whereas increasing the number of clusters separated cluster 3 into two separate clusters (each new cluster emphasizing the -1 and -2 nucleosomes, respectively, which are shown together in the current cluster 3). Therefore, we determined that four clusters showed the most distinct nucleosome architectures for primary LAC patient samples. We observed that across the genome there are differences between the normal and tumor patient samples, with the most pronounced changes occurring in clusters 1 and 4. We show a global loss of nucleosome occupancy and increased nucleosome phasing in the tumor sample. Clusters 1 and 4 display an impressive loss of nucleosomal occupancy upstream of the -1 nucleosome. These clusters indicate that changes in nucleosome occupancy in the grade one patients may play a role in the concerted gene regulation associated with transformation.

We next wanted to determine whether the 1,804 TSSs with altered nucleosomal structure shared in common between low-grade patients grouped into any particular cluster. We found that the majority (76%) of the 1804 shared genes were located in clusters 1 (32%) and 4 (44%) (582 and 799, respectively). Upon testing whether genes in each cluster were enriched for any particular gene ontology (GO) process, we found that each cluster had statistically significant GO enrichment (Fig. 2E) ^28^. Interestingly, genes in clusters 1 and 4 were each enriched for GO processes including chromatin and cancer-associated processes, such as nucleosome assembly, mRNA process, and mitotic G2 DNA damage checkpoint. Additionally, the genes in clusters 1 and 4 shared identical GO function for G-protein coupled receptor activity (P-value=1.51 x 10^-19^). Clusters 2 and 3 were enriched for genes with very general cell and molecular processes.

### Nucleosome distribution alterations are consistent between patients with early LAC

The similarity in the degree of difference between normal and tumor tissue for the grade one patients, the high overlap between patients for loci with altered nucleosome distribution, and the enrichment of those loci in related ontological categories indicated consistency between patients. We next visually inspected the nucleosome redistributions at specific loci to see whether the nucleosome distribution patterns at individual loci were similar between patients. The average nucleosome distribution plots for the 1,804 shared grade one genes showed many changes in nucleosome distribution among the grade one patients, and few changes among the grade three patients (Fig. 3A). The loss of occupancy in the grade one tumor is particularly clear in these average plots, with a majority of this loss occurring downstream of the TSS. Figure 3 shows five representative genes that are misregulated in adenocarcinoma: *ATM, CASC1, CDKL2, CCR10,* and *HKR1* (Fig. 3B-F) ^29^^-^^39^. In each case, the locus showed substantial differences between the grade-one patient normal and tumor samples. In a majority of cases we found that specific nucleosome distribution changes were consistent between and unique to the grade-one tumors (Fig. 3B-F, “Grade One” column, highlighted). Grade-three tumor samples rarely deviated from the nucleosome distribution pattern seen in normal tissue (Fig. 3B-F, “Grade Three” column, gray shaded). These results confirm a common mechanism driving the nucleosome distribution changes in the grade one tumors.

**Figure 3.**
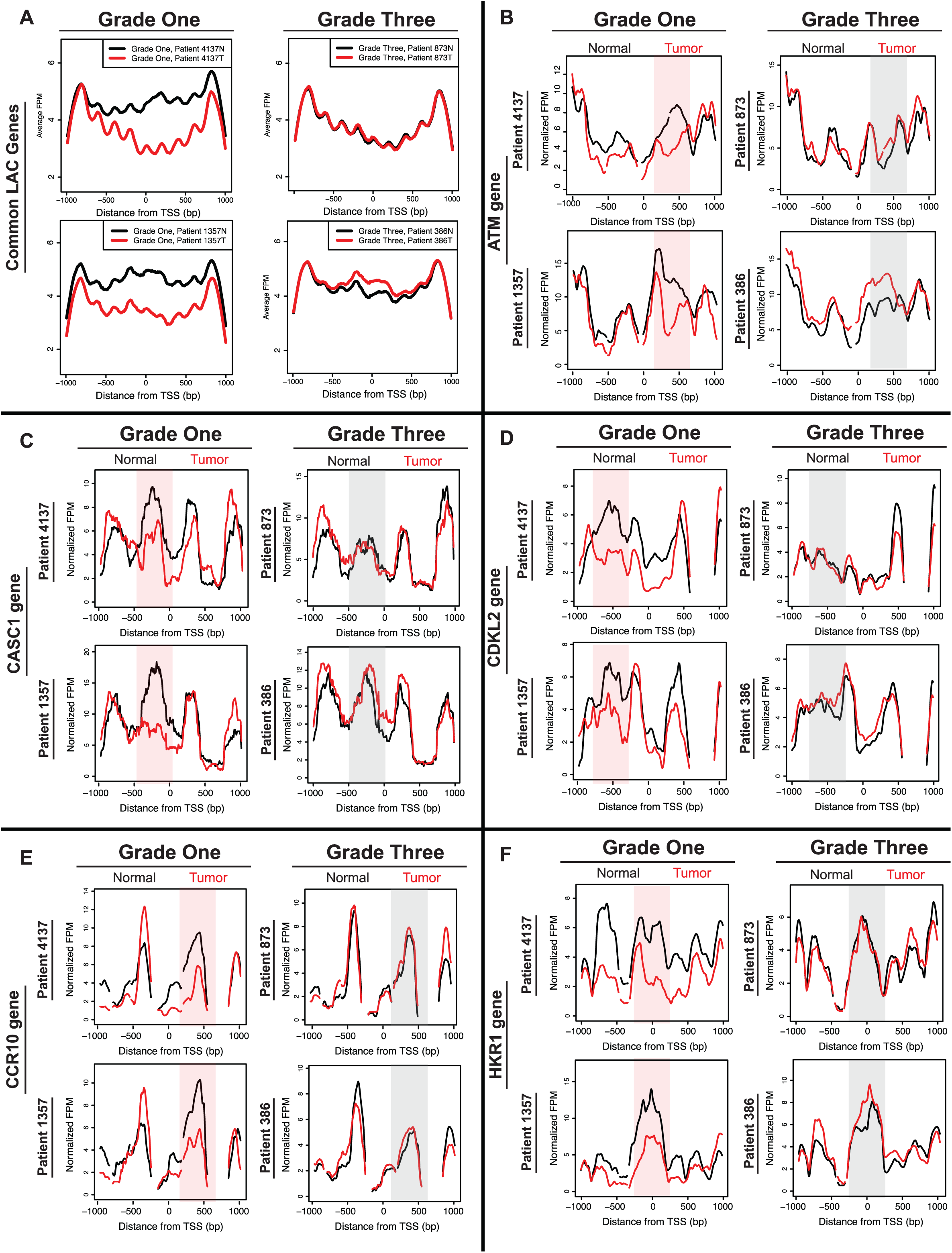
**Average and gene-specific plots show many nucleosome distribution changes that are consistent between patients in low-grade tumors and minimal changes in high-grade tumors.** (A) Average nucleosome distribution of normal tissue (black) and tumor tissue (red) for grade one patients (#4137 and #1357) and grade three patients (#873 and #386) for 1,804 genes shared between grade one patients. Five additional genes implicated in cancer are shown (B) ATM (C) CASC1 (D) CDKL2 (E) CCR10 and (F) HKR1. The x-axis represents a 2kb range of genomic position centered on the TSS, and the y-axis is fragments per million. Regions with most significant change in the grade one patients are highlighted in a shaded red for emphasis, while corresponding regions of no change in grade three patients are shaded in grey.

### Nucleosome distribution changes are driven by DNA sequence

Given the commonalities between the nucleosome distribution changes between the patients, we wanted to understand the influences driving the nucleosome distribution changes in the grade one samples. Nucleosome distributions are governed by the interplay between regulatory complexes, such as transcription factors and chromatin remodelers, and features intrinsic to the DNA sequence. We were interested in determining the extent to which DNA sequence contributed to the grade-one changes. We compared our experimentally determined nucleosome distributions to computationally predicted nucleosome occupancy scores based solely upon primary sequence ^40^^,^^41^. We reasoned that if DNA sequence played a role in in these distributions, then the predictions based upon the computational model would match the measured nucleosome distributions in the grade-one samples.

Of the 1,804 loci with nucleosome distribution changes shared between the grade-one patients, an average of ∼1,500 genes (85%) had a higher correlation with the DNA-encoded nucleosome positions than the matched normal sample, indicating that those loci are moving to positions favored by the underlying DNA sequence (Fig. 4A). In contrast, the same loci in the grade three tumors did not show increased agreement with the DNA-encoded positions; rather, the correlations between the grade three tumor and model based on DNA sequence were similar to those between the matched normal and model. Overall, the loci for the average grade one tumor data showed an average 215% increase in correlation with the computational model, as compared with the average normal data. The grade three tumors showed a 17% decrease in correlation to the computational model compared to normal.

**Figure 4.**
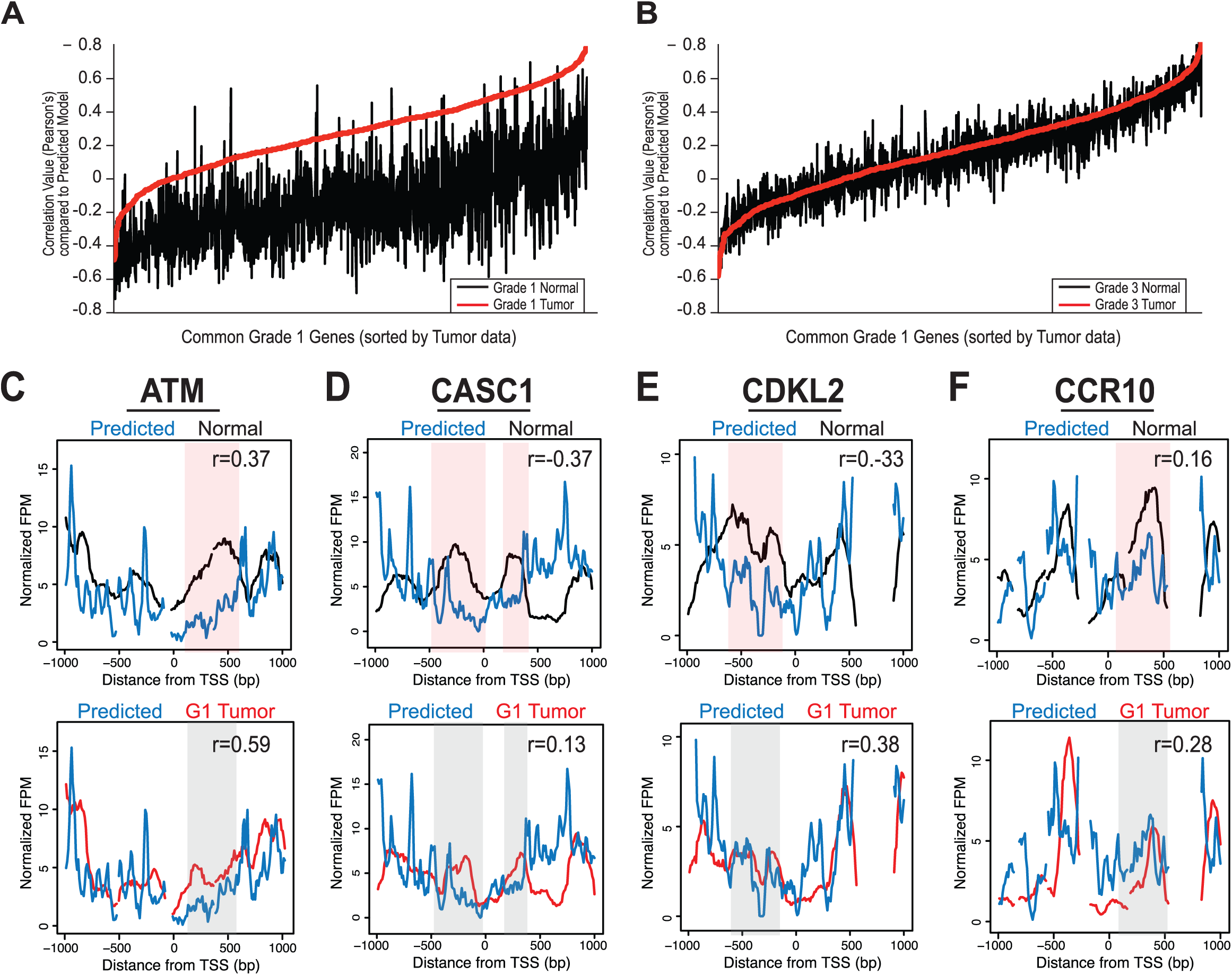
**Nucleosome distribution changes are driven by DNA sequence.** (A) Correlation values for the normal patient data (black) and grade one tumor patient data (red) versus the computationally predicted DNA encoded nucleosome occupancy model. The x-axis is the 1,804 genes in common between grade one patients, sorted on the correlation of the tumor data with the DNA encoded nucleosome occupancy model, and the y-axis is the Pearson’s correlation coefficient from the comparison of each data set versus the computational model values. (B) The correlation values for the average data from the normal tissue (black) and grade three tumor tissue (red) versus the DNA encoded nucleosome occupancy. Axes are identical to those in (A). The nucleosome distribution data for normal tissue (Normal, black lines) and grade one tumor tissue (red lines) are shown compared to DNA encoded nucleosome occupancy model scores (blue-DNA encoded, four genes from Fig. 3: (C) ATM, (D) CASC1, (E) CDKL2, and (F) CCR10. The x-axis represents a 2kb range of genomic position centered on TSS. The y-axis is the normalized fragments per million. Regions with most significant difference in the normal compared to the model are highlighted in a shaded red for emphasis, while corresponding regions of no change in the grade one tumor compared to the model are shaded in grey. Correlation values between the data and model are included for each gene; in all cases, the model is more highly correlated with the grade one tumors than the normal tissue or grade three tumors.

We then determined the DNA-directed nature of grade one tumor nucleosome distribution changes at individual loci. We co-plotted the nucleosome distribution data with the DNA-based model of nucleosome occupancy at the representative loci analyzed earlier. The agreement between the grade one nucleosome redistributions and the positions directed by the underlying DNA sequence was evident when we plotted the measured and predicted nucleosome distributions at specific loci (Fig. 4C-F). The correlation coefficient was always highest in the predicted versus grade-one data. These results suggest that DNA-encoded nucleosome signals direct nucleosomes to default positions upon transformation in early LAC.

### Altered nucleosome distribution in LAC potentiates transcription factor binding

In order to determine whether transcription factor binding occurred in the context of nucleosome redistributions in LAC, we first calculated regions of difference between normal and tumor throughout all TSSs for grade one and grade three patients. We found ∼18,000 regions of difference in the grade one and ∼6,000 regions of difference in the grade three samples by this method (Fig. 5A). The threshold applied to determine regions of difference was the most stringent cut-off that discriminated between the samples, while revealing a substantial enough number of regions to perform downstream analyses in the grade three patients since there were far fewer regions of difference than in the grade one patients. Overall, the total difference values for the grade one patients have a much higher range than the values for grade three patients. Therefore, although a region was determined above a threshold, the difference value was reliably lower in the grade three compared to the grade one patients, agreeing with our earlier observations that changes in nucleosome distribution occur early in the progression of cancer (Supplementary Figure 2A).

**Figure 5.**
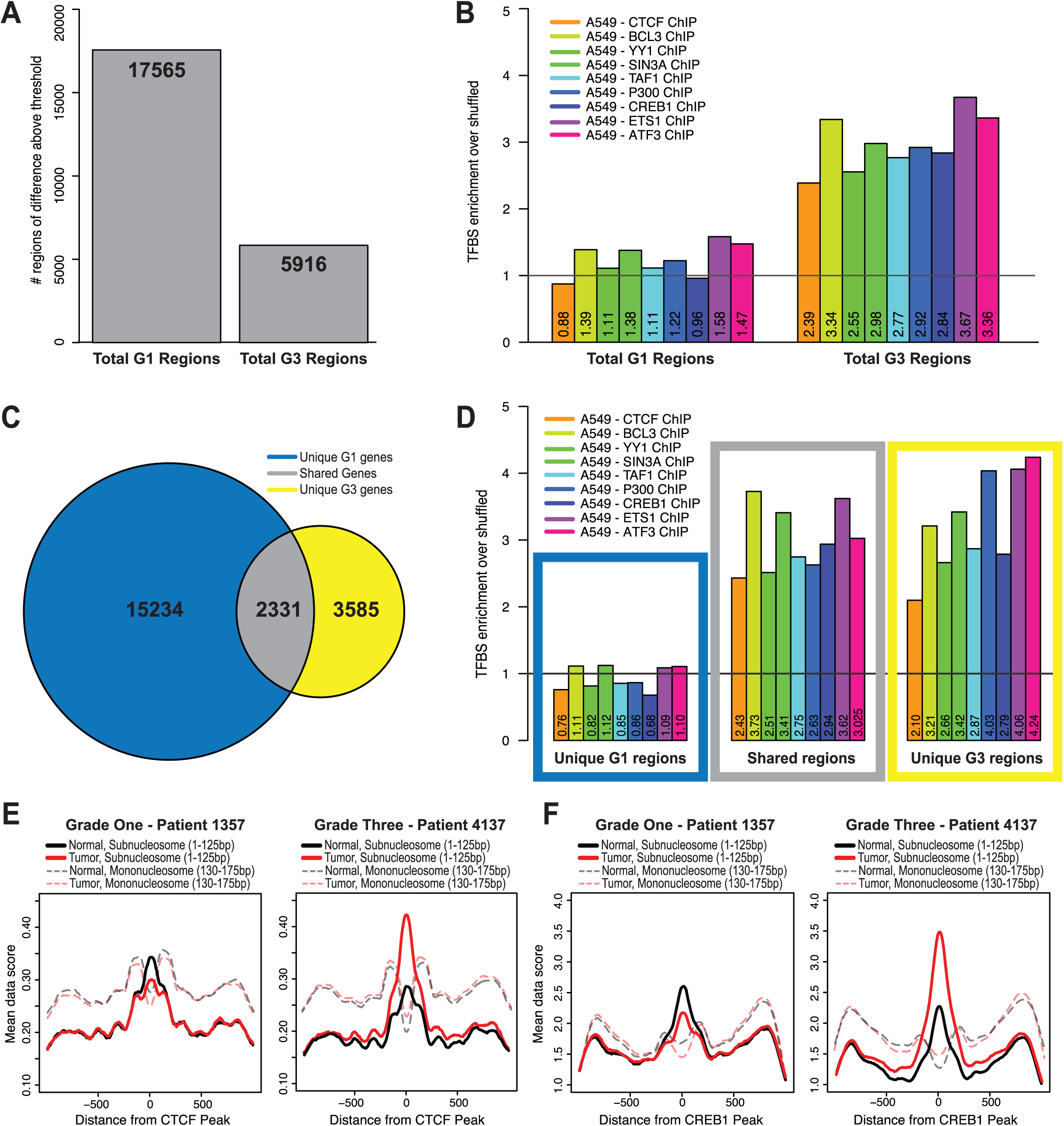
**Altered nucleosome distribution in LAC potentiates transcription factor binding.** (A) Regions of difference in grade one patients (17,565) and grade three patients (5,916), obtained by thresholding regions with a difference greater than 5. The threshold applied to determine regions of change was the most stringent cut-off that discriminated between the samples, while revealing a substantial enough number of regions to perform downstream analyses in the grade three patients since there were far less regions of change than in the grade one patients. Regions common between patients were merged so that duplicated regions were removed. We identified 17,565 regions of difference in the grade one patients and 5,916 regions in the grade three patients. (B) We determined the number of transcription factor binding sites for each region of difference in the grade one and the grade three patients, and calculated the ratio between the observed/shuffled. The black vertical line is drawn at 1, and values above 1 represent enrichment and below 1 represent depletion of transcription factor binding sites in the regions. We performed this analysis for nine transcription factors: Ctcf (GSM803456), Bcl3 (GSM1010775), Yy1 (GSM1010794), Sin3a (GSM1010882), Taf1 (GSM1010812), P300 (GSM1010827), Creb1 (GSM1010719), Ets1 (GSM1010829) and Atf3 (GSM1010789). (C) Venn diagram of overlap between regions corresponding to genes between the grade one patients and grade three patients: regions unique to grade one patients, shared regions, and regions unique to grade three patients. (D) For each of the categories from panel C, we determined the enrichment of transcription factor binding sites at the regions of difference through the same procedure and for the same nine transcription factors from panel B. The colored boxes correspond to the categories determined from the Venn diagram in panel C. Subnucleosomal fragment data (<125bp) for normal (black lines) and tumor (red lines) for each grade one patient #1357 and grade three patient #873 were aligned and centered on representative transcription factor data (E) Ctcf (GSM803456) and (F) Creb1 (GSM1010719) peaks from ChIP-seq in A549 cells. Additionally, nucleosome size fragment scores (130-175bp) for normal (shaded, dashed black lines) and tumor (shaded, dashed red lines) for each patient were also aligned and centered on Ctcf and Creb1 peaks.

Using transcription factor binding site (TFBS) data identified by ChIP-seq in a lung adenocarcinoma cell line (A549) we quantified binding sites for nine transcription factors Ctcf (GSM803456), Bcl3 (GSM1010775), Yy1 (GSM1010794), Sin3a (GSM1010882), Taf1 (GSM1010812), P300 (GSM1010827), Creb1 (GSM1010719), Ets1 (GSM1010829) and Atf3 (GSM1010789) at the regions of difference in grade one and grade three patients ^42^. In order to determine enrichment, we shuffled the TFBSs identified in the A549 study and then calculated a ratio of the number of binding events in the regions of difference to that shuffled control (a value of one indicates no significant enrichment or depletion compared to the shuffled data). We found significant enrichment over shuffled TFBSs tested at regions of difference in the grade three patients, and depletion of TFBSs in the grade one patients (Fig. 5B). In order to verify that the TFBS depletion in the regions of difference was a feature exclusive to the grade one patients, we first determined the overlap of regions of difference between the grade one and grade three patients, and we found that 2,331 regions were shared in common (Fig. 5C, Supplementary Figure 2B-D). When we compared each of these categories to TFBSs, we found that the regions unique to grade one patients were depleted of TFBSs, whereas the shared genes and the genes unique to grade three patients were highly enriched for TFBSs (Fig. 5D). These results suggest that changes in nucleosome distribution in the grade one tumors broadly alter access to the genome, and the changes that persist in the grade three patients are likely the result of differential transcription factor binding.

To investigate whether the nucleosome distribution changes in the grade one tumors exposed DNA at loci for genomic licensing, we measured the proportion of subnucleosomal MNase protected fragments at regulatory factor binding sites. It has been shown that subnucleosomal fragments (< 100±20 bp) derived from MNase digestion of DNA may act as a proxy for protection by DNA-binding proteins, such as transcription factors.^11^^,^^13^ To test the hypothesis that nucleosome distribution changes alter access to the genome potentiating transcription factor binding, we examined binding alterations at specific transcription factor binding sites. Using subnucleosomal fragment data for all fragments less than 125bp from grade one and grade three patients we plotted all reads centered on the binding sites for Ctcf (Fig. 5E) and Creb1 (Fig. 5F). We found that when normal and tumor tissue were compared, there is a 13% decrease in binding of Ctcf and a 19% decrease of Creb1 binding in the grade one patients, while for the grade three patients there was a 47% increase in binding of Ctcf and a 31% increase of Creb1 binding. We also plotted the nucleosome size fragments with a size range of 130-175bp for each patient and transcription factor, and confirmed that these inferred transcription factor binding events were associated with nucleosome free regions, typical of regulatory factor binding sites ^11^^,^^13^. Taken together, our results suggest that nucleosome redistribution, which provides the opportunity for transcription factors to bind with a greater probability, is a potentiating event in the progression of cancer.

### Nucleosome distribution changes are widespread in the progression of CRC, consistent with LAC, driven by DNA sequence and potentiate transcription factor binding

To determine whether the widespread nucleosome redistributions were a feature unique to LAC or if nucleosome alterations are a common characteristic of adenocarcinoma types, we mapped the nucleosome distribution in CRC patients. We performed mTSS-seq on matched normal tissue and tumors of stage two (S2), stage three (S3) and stage four (S4). We calculated the correlation between normal and tumor nucleosome distribution for each patient, and found widespread changes in patients with early-CRC (S2 and S3). There were 2,133 genes shared in common between these patients early CRC patients. We next compared these 2,133 common CRC genes with the 1,804 common LAC genes, and found 709 genes with altered nucleosome distribution shared between LAC and CRC. The nucleosome distribution at the ATM, HKR1, NOP16, and KIF2B genes for early LAC and all CRC patients showed that the nucleosome redistributions identified are consistent between the early CRC CRC patients, and are absent in the advanced (S4) CRC patient (Fig. 6A-D). These plots also show that nucleosome redistributions in early CRC are consistent with changes in early LAC patients for ATM, HKR1, NOP16, and KIF2B genes (Fig. 6A-D) ^29^^,^^35^^,^^38^^,^^39^. This consistency between the two early adenocarcinomas is an important finding, indicating that nucleosome redistributions are a common genomic feature of early transformation.

**Figure 6.**
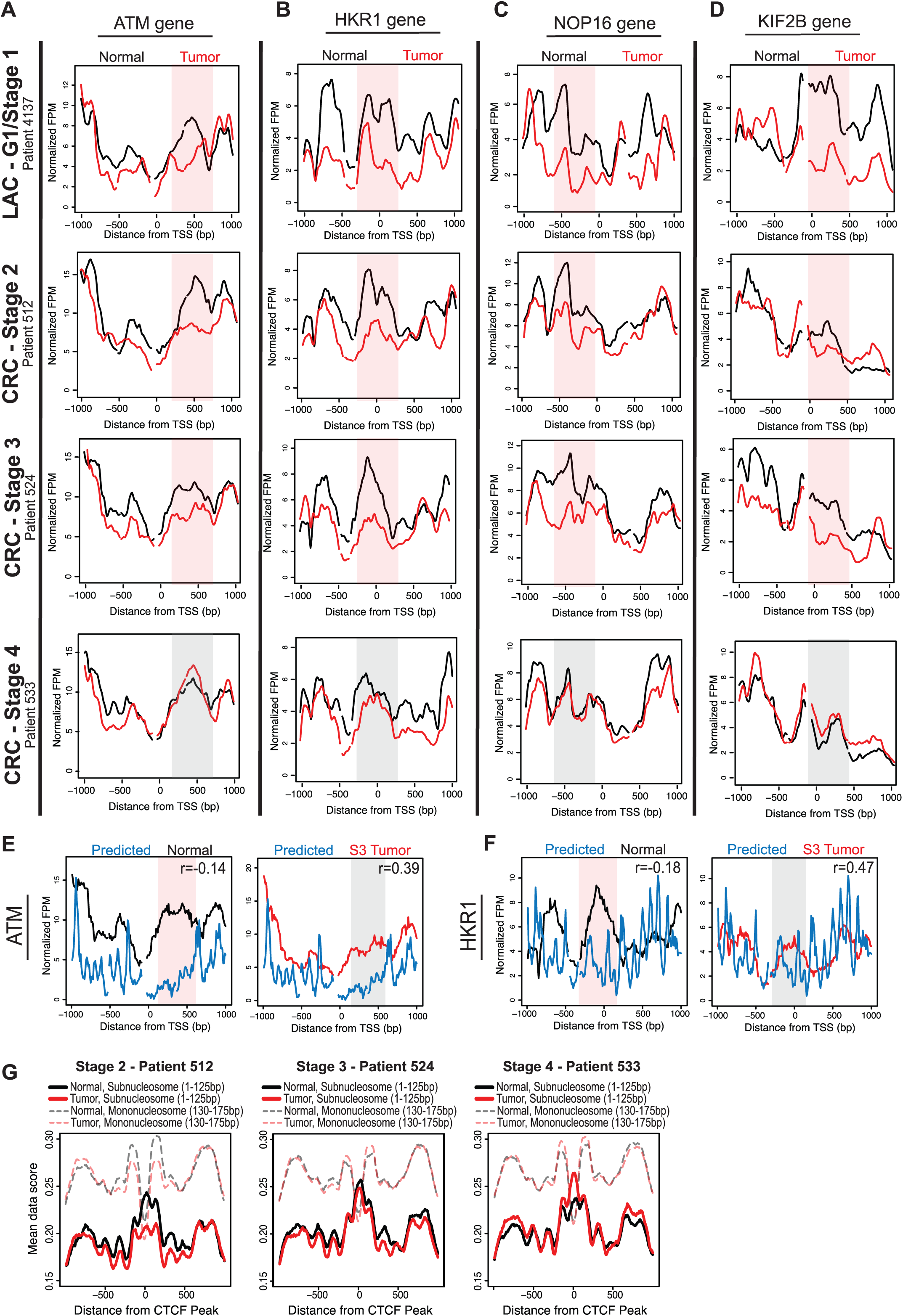
**Nucleosome distribution changes in early-CRC are widespread, concordant with LAC changes, DNA-directed and potentiate transcription factor binding.** (A) Nucleosome distribution plots at the ATM gene are shown for normal (black lines) compared to matched grade one LAC tumor (as seen in Fig. 3B; red lines), and for the normal compared with matched S2, S3 and S4 CRC tumors (red lines). The x-axis represents the TSS +/- 1kb, and the y-axis is fragments per million. Regions with most significant change in the grade one patients are shaded in red for emphasis, while corresponding regions in grade three, which are unchanged between normal and tumor samples, are shaded in grey. Three other cancerrelated genes are shown to illustrate the widespread nucleosome distribution changes in the progression of CRC, and concordance with LAC changes (B) HKR1 (C) NOP16 (D) KIF2B. The nucleosome distribution data for normal tissue (black lines) and S3 CRC tumor tissue (red lines) are shown compared to predicted nucleosome occupancy based on DNA sequence (blue lines), for genes from Fig. 4C and this figure B: (E) ATM, and (F) HKR1. The x-axis represents the TSS +/- 1 kb, and the y-axis is the normalized fragments per million. Regions with most significant difference in the normal compared to the model are shaded red for emphasis, while corresponding regions in the S3 tumor are shaded in grey. Additionally, correlation values between the data and model are included for each gene; in all cases, the model is more highly correlated with the S3 tumors than the normal tissue. (G) Subnucleosomal fragment data (<125bp) for normal (black lines) and tumor (red lines) for each CRC patient were aligned and as a representative centered on transcription factor data for Ctcf (GSM803456) peaks from ChIP-seq in A549 cells. Additionally, nucleosome size fragment scores (130-175bp) for normal (shaded, dashed black lines) and tumor (shaded, dashed red lines) for each patient were also aligned and centered on Ctcf peaks.

To assess the role of *cis* and *trans*-acting factors governing nucleosome redistributions in the progression of CRC, we first compared the experimentally determined nucleosome distributions for the common CRC genes to the computationally predicted model. We found that the early CRC tumors had a higher correlation than normal with the predicted model at over 58% of genes. The S3 CRC tumor and matched normal data compared to the predicted model at ATM and HKR1 genes showed a greater agreement between the predicted model and the tumor than between predicted model and the normal data (Fig. 6E-F). Finally, centering subnucleosomal fragments on CTCF binding peaks showed a decrease in binding in the early CRC patients and an increase in binding in the more advanced CRC patients (Fig.6G). These results confirm that nucleosome distribution changes are widespread and consistent in both early LAC and CRC, are directed by the underlying DNA sequence, and likely potentiate an increase in transcription factor binding in advanced cancer.

## DISCUSSION

A full comprehension of the relationship between chromatin structure and genome function in cancer necessitates genome-wide chromatin structural measurements at multiple points in time throughout cancer progression. Although there have recently been a handful of extremely important studies measuring nucleosome distribution in a variety of organisms, there have been no genome-wide nucleosome distribution maps in primary patient tumors compared to their matched normal tissue ^5^^-^^7^,^9^,^11^,^12^,^27^,^41^,^43^. To meet this need, we developed an innovative approach, mTSS-seq, to comprehensively measure genome wide nucleosome distribution changes in the progression of cancer. In this study, we validated our approach for high resolution, genome-wide nucleosome distribution mapping utilizing data from a very high quality human MNase-seq nucleosome mapping study, and our previous microarray based nucleosome maps from LAC patients. We anticipate that this first comprehensive analysis of the relationship between chromatin structure and genome regulation in the progression of cancer will pave the way for similar detailed studies in other diseases.

In our previous work we introduced a model in which we identified widespread changes in nucleosome distribution as a feature specific to low grade cancer. We derived that model from the study of ∼900 cell cycle- and immunity-related genes ^9^. Because the original model was based on a limited set of genes, we wanted to determine whether the changes in nucleosome distribution were a widespread feature across all genes in the human genome. Therefore, we developed the mTSS-seq target enrichment platform to test and expand our original model across the entire human genome in multiple patient samples. Using mTSS-seq, we measured changes in nucleosome distribution between tumor and normal tissue, for each LAC patient, and made three initial striking discoveries: 1) nucleosome distribution changes are indeed a widespread feature across the entire genome in the tumor samples from early LAC patients, suggesting global dysregulation of chromatin remodeling as an early transformation event; 2) nucleosome distribution changes are consistent among the early LAC patients, suggesting a common dysregulation among patients; and, 3) widespread nucleosome distribution changes are comparatively absent in more advanced tumors, suggesting that the remodeling dysregulation does not persist into advanced tumors. Widespread nucleosome distribution changes that appear in low-grade as opposed to more advanced tumors that are consistent between patients indicates an early, concerted genomic event in the progression of cancer. It is tempting to speculate that if changes in nucleosome distribution act as an indicator of impending transcriptional regulation, then our nucleosome distribution measurements could act as predictive indicators of early transformation events. This explanation is manifested in a recent report from our lab in which widespread, transient, DNA-directed nucleosome redistributions were observed at immune loci upon reactivation of Kaposi’s sarcoma-associated herpesvirus (KSHV), an oncogenic viral system ^41^.

In the current study, we have additionally expanded the original model by demonstrating that the nucleosome distribution changes occur through genetically-encoded regulatory signals: the nucleosomes in the grade one tumors are remodeled to positions encoded by the DNA sequence. Again, this observation is consistent with our work on KSHV, in which we established that transient nucleosome redistributions, rather than basal architectures, adopt locations favored by the underlying DNA sequence ^41^. In the current study, we demonstrate that the low-grade tumor samples had a higher correlation with the predicted model as compared to normal tissue at over 85% of remodeled genes, indicating that nucleosome distribution alterations are driven by the underlying DNA sequence ^18^^,^^40^^,^^41^. An appealing interpretation that reflects these consistent grade-one nucleosome distribution alterations is that the redistributions result from the misregulation of a chromatin remodeling complex that culminates in nucleosomal redistribution to DNA-directed positions. This is conceivable given the evidence in the literature on genomic dysregulation through mutation of chromatin remodeling complexes in cancer determined by exome sequencing ^44^^-^^48^. An intriguing, remaining question centers on the apparently ephemeral nature of these grade-one changes, and the degree to which redundant and overlapping chromatin regulatory activities play a role in the complex progression of cancer.

To answer questions regarding the effects of these apparently transient nucleosome redistributions, we provide evidence that nucleosome redistributions likely potentiate transcription factor binding events. Using subnuclesomal sized DNA fragments as an indicator of transcription factor binding, we measured depletion or enrichment of transcription factor sized protections at known transcription factor binding sites identified by ChIP in A549 lung cancer cells. We observed an increase in the presence of subnucleosomal fragments in high grade tumors compared to normal tissue at known transcription factor binding sites identified by ChIP in A549 lung cancer cells, indicating the presence of a sequence-specific DNA-binding protein ^42^. This increase in transcription factor binding in advanced tumors relative to the normal tissue and grade one tumors suggests that nucleosome redistributions early in the progression of cancer potentiated the licensing of these regulatory factors.

A particularly remarkable extension of the original study is the finding of widespread nucleosome redistributions in the progression of CRC that are concordant with the changes observed in LAC. In LAC, nucleosome distribution alterations are widespread in low grade tumors (grade one, stage one), and these alterations are not seen in high grade tumors (grade three, stage two). We showed that these widespread nucleosome distribution alterations also occur in early CRC (stage two and three), and these changes are relatively absent in more advanced CRC (stage four). There is a high overlap of genes with nucleosome distribution alterations between LAC and CRC. Moreover, we have shown that the redistributions in CRC have a strong agreement with genetically encoded nucleosome distribution signals, indicating that the nucleosome distribution changes are DNA-directed as in LAC. The discovery of increased transcription factor binding events in advanced tumors was also observed in CRC patients. Utilizing a high-resolution, genome-wide technology to identify widespread chromatin structural changes in early tumors across multiple cancer types while defining the functional regulation through analysis of *cis-* and *trans-* acting factors validates the power of this approach to study chromatin structure in the progression of multiple cancers and disease states.

Taken together these results clarify structure-function relationships in the human genome, and support a hierarchical mechanism for chromatin mediated genomic regulation ^41^. This study demonstrates that widespread, DNA-directed nucleosome redistributions are limited to early tumors in LAC and CRC. This hierarchical model describes the interpretation that these nucleosome redistributions likely allow for inappropriate regulatory licensing in cancer (Fig. 7A). Indeed, inappropriate genomic licensing is frequently cited as a characteristic of transformed phenotype ^49^^,^^50^. We propose that in the later stage and grade tumors when nucleosomes return to their basal positions, the regulatory machinery is altered, and contributes to the progression of the disease (Fig. 7B). This comprehensive and integrated analysis of the relationship between chromatin structure and the progression of cancer has allowed us to define nucleosome alterations as generally exploited sites of concerted dysregulation in cancer.

**Figure 7.**
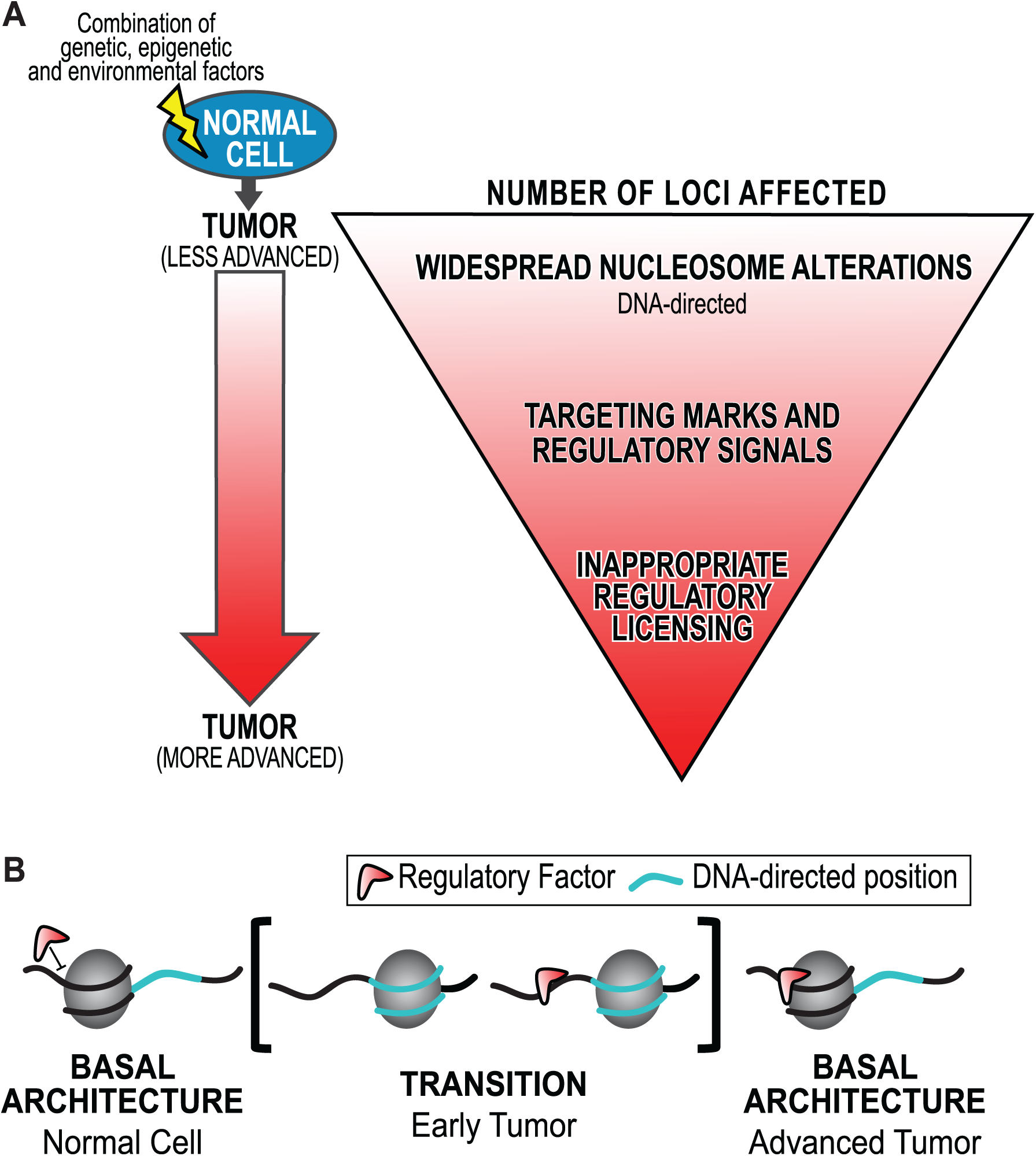
**A model for chromatin based hierarchical genome regulation.** (A) We have developed a model describing chromatin-based hierarchical genome regulation. In this model, a superset of genomic loci is made available for licensing through transient DNA-directed nucleosome redistributions: a genomic intermediate. Loci in a physiology with the appropriate regulatory machinery will be licensed for a genomic response. Those without the regulatory machinery will not be affected. This model maximizes the potential for multiple concerted responses with a limited number of genomic architectures. However, if any point of this hierarchy is disrupted oncogenic transformation can occur. (B) An interpretation of these results is that regulatory factors are initially unable to bind basal nucleosome architecture (as in the normal). Inappropriate widespread nucleosome redistributions to DNA-directed positions in early tumors potentiate the binding of regulatory factors (such as transcription factors) outside of their physiological context. Nucleosomes return to their basal architecture in advanced tumors, possibly through redundant and compensatory remodeling machinery. However, the regulatory mark remains, which further contributes to the progression of cancer.

### Materials and Methods

#### Patient samples and tissue processing

Primary samples from surgically removed tumors of lung adenocarcinoma patients, and corresponding normal tissue were obtained from the University of Massachusetts Medical School (UMMS) Tissue Bank, and prepared as previously described ^9^. Primary samples from colorectal adenocarcinoma patients with a surgically removed tumor, and corresponding normal tissue were obtained from the Mayo Clinic through Dr. Lisa Boardman. A total of 7 tumor specimens were included in this study (LAC: two grade one, two grade three; CRC: one of each stage one, two and three), with matched normal tissue for each tumor specimen, for a total of 14 genomes that were sequenced. The tumor and normal material was snap-frozen in liquid nitrogen within 1 h after surgery. Samples were examined by board-certified pathologists, using hematoxylin and eosin staining. Samples were selected by grade and stage, and only samples with 80% or more tumor cells were included, as assessed by histological examination. Patient samples were anonymized, and we received patient history along with the samples. Harvesting of nuclei, MNase digestion and mononucleosomal isolation were performed on each sample as previously described ^9^. See Supplementary Table 1 for preparation information on each patient sample.

#### Mononucleosome DNA Library Preparation

MNase digested DNA sequencing libraries were prepared using the NEBNext® Ultra^TM^ DNA Library Prep Kit for Illumina® (NEB #E7370S/L), starting with thirty nanograms of input mononucleosomal DNA. Following end prep and adaptor ligation, libraries were cleaned-up with AMPure® XP Beads (Beckman Coulter, Inc. #A63881) without size selection due to the original input of a size population of ∼150bp. Universal and indexed sequences were added through 8 cycles of PCR, using NEBNext® Multiplex Oligos for Illumina® (Index Primers Set 1, NEB #E7335S/L). The NEBNext® Multiplex Oligos kit contains indices 1-12 which correspond to the identical product if using Illumina® TruSeq primers. The libraries were quantity and quality checked using the Qubit Fluorometer High Sensitivity Kit and Agilent High Sensitivity DNA kit on the Agilent 2100 Bioanalyzer. The average size of material across all libraries was 275bp, and the average total material in this region was more than 90%; there were no adapter or primer dimers.

#### Solution-based Sequence Capture, enabling TSS-enrichment

We used a custom designed Roche Nimblegen SeqCap EZ Library SR to capture ∼2kb regions flanking the TSS for every gene in the human genome, using the HG19 build. The TSS sequences were repeat masked, so only unique probes were included. We performed the sequence capture according to the manufacturer’s protocol. Following a 72 hour capture hybridization, we performed a 15 cycle PCR amplification using the TruSeq primers 1 (AAT GAT ACG GCG ACC ACC GAG A) and 2 (CAA GCA GAA GAC GGC ATA CGA G). We then performed a quantitative real-time PCR to confirm that regions within the sequence capture were successfully enriched, and that regions excluded from the capture were depleted post-capture. We selected three regions within the 2kb TSS of genes that we knew to be in the SeqCap design (on-target), and for the same three genes we selected regions outside of the 2kb TSS that were not in the SeqCap design (off-target). The on-target and off-target regions and primer sequences can be found in Supplementary Table 2. Dilutions were made in elution buffer to 10 nM stock in 0.05% Tween-20.

#### Illumina Flowcell hybridization and sequencing

The multiplexed samples were loaded at 12pM on two lanes of an Illumina HiSeq 2500 system, HiSeq Flow Cell v3. For the HiSeq, the suggested range is 10-20pM. Kits used were the TruSeq PE Cluster Kit v3-cBot – HS and the TruSeq SBS Kit v3.

There are two measures for data quality: the first is clusters that pass filter (PF) and the other is a quality score, which is given as a percentage of reads > Q30. The reads are based on the reads that pass the chastity filter not the Q30 filter. In addition, each lane was spiked with 1% PhiX as the control. The software performs real-time reporting of error rates for the PhiX spike-in lanes. The sequencing was a paired-end 50 bp run on the HiSeq, using HiSeq Control Software (HCS) version 2.0. The LAC lane had cluster density of 695K/mm[2], a PF of 94%, and 96.6% of the reads having a quality score >Q30. The CRC lane had cluster density of 736K/mm[2], a PF of 94%, and 96.1% of the reads having a quality score >Q30. The samples that were sequenced by on the MiSeq were run on 3 lanes, and was paired-end 150bp sequenced (Supplementary Table 1 – sequencing processing). The first lane was loaded at 8pM and generated 1468 clusters k/mm2. The other two lanes were loaded at 4pM and obtained 681 k/mm2 and 658 k/mm2 clusters, respectively. MiSeq V2 reagents were used and the MiSeq default settings were applied to generate fastq files that contain only PF reads (pass filter). The reads were demultiplexed on the MiSeq using the default settings. Paired sequence reads have been deposited at http://bio.fsu.edu/∼broberts/NatureGenetics_data/.

#### Alignment and data processing bioinformatics

Casava software was used to demultiplex the indices in each lane. Illumina adapters were clipped from reads with cutadapt ^51^ and aligned to the hg19 human genome assembly with bowtie2 2.1.0 with default parameters ^52^. Unpaired and non-uniquely-mapped reads were discarded with samtools ^53^. Individual nucleosome footprints were extracted from BAM files with bedtools 2.17 ^54^. Nucleosome occupancy profiles were obtained by calculating the fragments per million that mapped at each base-pair in the probed regions with bedtools. Nucleosome dyad frequencies (midpoints) were obtained by calculating the sum of nucleosome dyads (fragment centers) in 100-bp windows at a 10-bp step-size with bedtools. Data were subsequently processed in R 2.15.1 ^55^. Data was uploaded to the UCSC genome browser for further analysis (<u>http://genome.ucsc.edu</u>) ^56^^,^^57^.

